# Food-associated calls in disc-winged bats

**DOI:** 10.1101/2024.02.25.581993

**Authors:** Gloriana Chaverri, Rachel A. Page

**Author notes:** Corresponding author: Gloriana Chaverri.

## Abstract

Animals that engage in social foraging can produce food-associated calls that elicit two main responses in receivers: the recruitment of other individuals to a foraging site, and an increase in feeding-related behaviors in conspecifics. Spix’s disc-winged bat, *Thyroptera tricolor*, is a highly gregarious species that lives in stable social groups and relies on group call-and-response vocalizations to find ephemeral roosting sites. Because this bat also is known to feed on resources that are abundant but ephemeral – for example insect swarms – we hypothesized that it likewise emits vocalizations that serve to recruit conspecifics to a foraging site and elicit food consumption. We found that indeed, feeding bats emitted distinct vocalizations exclusively while consuming an abundant prey item, a call type that has not been previously described in the acoustic repertoire of any bat to date. We also observed that these “food calls” prompted responses typically associated with food-calling: they increased both feeding-related behaviors and social recruitment. Specifically, we observed that the onset of the consumption of novel prey items was strongly associated with the emission of food calls but not with other types of sounds. In addition, individuals approached a speaker broadcasting food calls, especially when food calls had not been broadcast before, while other types of sounds did not consistently prompt inspection. Taken together, these results suggest that *T. tricolor* coordinates foraging behavior through the emission of food-associated communication calls.

**SUMMARY STATEMENT:** This study provides evidence of food-associated calling in Spix’s disc-winged bats (*Thyroptera tricolor*), revealing its association with feeding contexts and its potential role in prompting social recruitment and feeding-related behaviors.

## INTRODUCTION

Social foraging occurs when interactions among individuals, either cooperative or exploitative, involve foraging (Giraldeau & Caraco, 2000). Organisms that engage in social foraging feed near other foragers. Close spatial proximity of individuals that consume similar food items, however, can result in significant costs given the depletion of available resources with increasing group size and aggressive interactions that may ensue (Grove, 2012; Johnson, 2004; Rose & Soole, 2020). Foraging in groups can also provide many benefits, however, including an increase in the sensory ability to detect patchily distributed prey (Cvikel et al., 2015; Dechmann et al., 2010; Kohles et al., 2022), a reduction of individual investment in vigilance (Elgar, 1989), a decrease in the per-capita risk of predation (Foster & Treherne, 1981), and the recruitment of conspecifics for the defense against potential predators and intruders (Cunha et al., 2017; Marzlufi & Heinrich, 1991). As expected given these advantages, social foraging has been observed across a wide range of taxa, including cliff swallows (*Hirundo pyrrhonota*; Brown, 1988), capuchin monkeys (*Cebus apella*; di Bitetti & Janson, 2001), and guppies (*Poecilia reticulata*; Day et al., 2001), among many others.

When individuals within foraging groups encounter profitable foraging patches, they often emit food-associated calls, or “food calls”. The emission of food calls is known to elicit two main responses in receivers: an increase in feeding-related behaviors, and the recruitment of other individuals to the foraging site (Clay et al., 2012). In domestic fowl (*Gallus gallus*), for example, males emit food calls when discovering food, and this causes rapid recruitment of females (Evans & Marler, 1994; Marler et al., 1986). Food calls have also been considered to be arousal stimuli that prompt feeding-related behaviors, such as searching (Evans & Evans, 2007), pecking (Wauters & Richard-Yris, 2002), and consuming provisioned food (Kitzmann & Caine, 2009). Most studies agree that the presumed function of these acoustic signals is to inform conspecifics of the location of shareable food sources (Clay et al., 2012; Giraldeau & Caraco, 2000). By recruiting group members to a feeding patch, individuals may gain some of the aforementioned benefits of social foraging, in addition to increasing inclusive fitness by aiding close kin locate a profitable food source (Hauser & Marler, 1993; Judd & Sherman, 1996), enhancing social status (Krunkelsven et al., 1996), or strengthening social bonds (Slocombe et al., 2010). The latter benefits may be particularly relevant for species that form socially stable groups (Clay et al., 2012).

A species in which we would predict both social foraging and food-associated calling is Spix’s disc-winged bat, *Thyroptera tricolor*. This species forms cohesive groups that are comprised of philopatric males and females, which are known to remain within their natal groups for several years and are thus closely related (Buchalski et al., 2014; Chaverri, 2010; Chaverri & Kunz, 2011). Disc-winged bats use highly ephemeral roosting resources that force individuals to locate a new site almost daily (Vonhof & Fenton, 2004). Despite these constant movements, individuals are able to maintain group cohesion by exchanging acoustic signals during flight and also when bats locate a new roost site (Chaverri & Gillam, 2016). When bats are flushed from a roost during the day, individuals constantly emit “inquiry calls”, which allow them to remain in close contact with other group members during flight. When one individual locates and enters a roost, it usually starts emitting “response calls” after hearing inquiry calls; flying bats then quickly enter this roost (Chaverri et al., 2010). Both types of acoustic signals have strong individual signatures, and bats seem to discriminate among the calls of group and non-group members, preferentially joining the former (Araya-Salas et al., 2020; Chaverri et al., 2013; Gillam & Chaverri, 2012). Inquiry calls are emitted by bats in flight not only during the day, but also at night (Montero & Gillam, 2015), suggesting that individuals actively maintain group cohesion not just in roost finding but also while foraging.

The diet of *T. tricolor* includes various arthropods, such as jumping spiders (Aranea) and insects, particularly hemipteran within the family Fulgoridae and various species of diurnal Diptera and wingless larval Lepidoptera and Hymenoptera, among others (Dechmann et al., 2006; Whitaker & Findley, 1980). Some of these insects may be found in large swarms, for example termites during their nuptial flights (Dechmann et al., 2006). The latter is relevant for understanding bat social interactions while foraging, given that clumped feeding resources can reduce the costs of sharing resources and can thus facilitate social foraging and food-calling (Clay et al., 2012; Giraldeau & Caraco, 2000). Notably, we have observed *T. tricolor* produce audible calls when feeding on mealworm larvae (*Tenebrio molitor*), which prompts consumption of provisioned food in conspecifics. Given what we know about *T. tricolor*’s social structure, its feeding habits, and the constant emission of acoustic signals in various social contexts, in this study we wanted to determine if individuals emit food calls. Specifically, we aimed to 1) describe the calls emitted while feeding and 2) test their effect on conspecifics. We hypothesized that, if the signals we recorded were indeed food calls, their playback would elicit consumption of provisioned food items and recruitment.

## METHODS

### Field methods

During the afternoon, we searched for groups of bats in Soberanía National Park near Gamboa, Panamá. Individuals were captured within their roosts, the developing tubular leaves of plants in the order Zingiberales. Upon capture, bats were placed in cloth bags. All individuals were marked with passive integrated transponders (Mini HPT8 Pit Tag, Biomark Inc., Idaho, United States). These markers have a unique numbering that allows individual identification of each animal with a digital reader (HPR Lite, Biomark). These tags are very small (1.4 x 8.5 mm) and weigh just 0.09 g (ca. 2 % of the bat’s body mass). All individuals were sexed and aged, and we registered the location of each group with a GPS (GPSMAP 64csx, Garmin).

### Recording food calls

In previous projects, we heard *T. tricolor* produce audible calls when hand-fed with mealworm larvae (*Tenebrio molitor*). To investigate these calls, in this project we held bats individually and recorded them as we hand-fed them mealworms. We made all recordings using a CM16 microphone connected to an UltraSoundGate Recorder (model 116 Hme, Avisoft) connected to a laptop computer running Avisoft Recorder software with 500 kHz sampling rate, and a resolution of 16 bits. Bats were recorded at 10 cm from the microphone, and the USG recorder’s gain was set at medium. Each bat was recorded three times, once in the late afternoon (between 16:00 – 17:00 hours), then again after sunset (around 18:00), and then once more after flying for ca. 15 min in a small flight cage. During each of the three recording sessions bats were fed between 2-4 mealworms. After the third trial bats were released at their capture site (see below).

### Playback files

We recorded a total of 15 individuals from 8 different social groups while feeding on mealworms. From these individuals we extracted recordings of chewing sounds from 4 bats, while high quality (i.e. those with high signal-to-noise ratio) food calls were collected from 6 individuals (Fig. 1, Supplementary Table 2). These food calls (n = 19) and chewing sounds (n=4) were then used to create sound files used in playback experiments. We created 5 different sets of files, each 60 sec long: 1) food calls only, 2) food calls + chewing sounds, 3) chewing sounds only, 4) pink noise, and 5) silence. All playback files were generated using a 500 kHz sampling rate and a resolution of 16 bits. During playback trials, the amplitude was set to similar levels as those originally recorded in the previous section. We created 10 playback files for each of the first 3 sets (food calls, food calls + chewing sounds, chewing sounds) for a total of 30 files; each file contained 5 different food calls and/or 3 chewing sound bouts from different individuals. We only created one pink noise file and one silence file, each 60 s long.

**Fig. 1.**
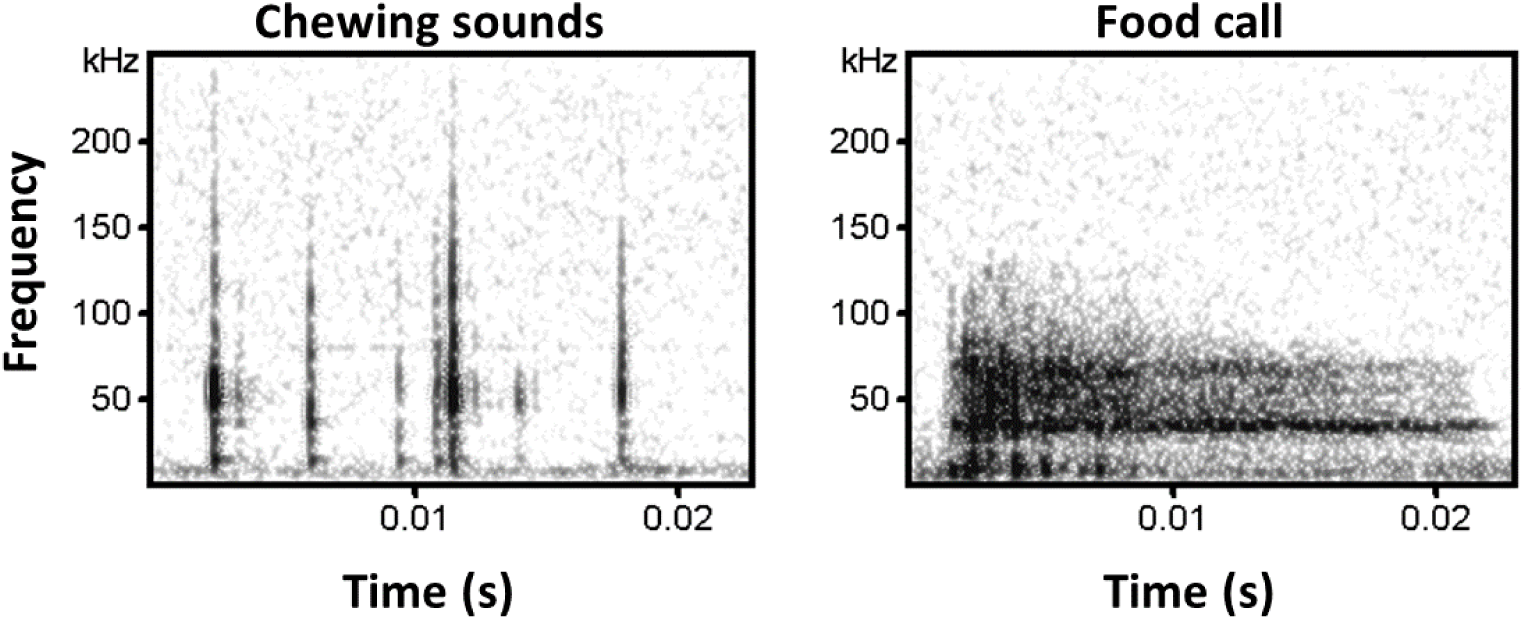
Spectrograms showing examples of chewing sounds (left) and food calls (right). Both sounds are produced by *Thyroptera tricolor* when eating mealworm larvae.

Playback files were generated as follows. First, we created 10 files with chewing sounds only. Each file had 5 bouts of chewing sounds, each 6 sec long (Fig. 2). For each playback file we randomly selected three 6-s bouts of chewing sounds from different individuals and combined them so that in the end we would have 2 sounds repeated twice, for a total of 5 in a single playback file. Each playback file then contained first a period of silence (2.5 s), then one 6-s chewing bout, then 6 sec of silence, then another 6-s chewing bout, and so on until completing 60 s (Fig. 2a). After the chewing-only playback files were created, we added the food calls to those same 10 files (Fig. 2b). Within each 6-s chewing bout we added 3 different food calls (from different individuals), each broadcast in a rapid succession of 5 calls (with a call interval of ca. 0.20 s ± 0.02 s, as recorded when hand-feeding bats), such that each 6-s bout had 15 food calls in total. For each of the other bouts we repeated the same process, using the same calls but in random order. Finally, we removed the chewing sounds from the files, leaving the food calls to create the food-call-only files (Fig. 2c).

**Fig. 2.**
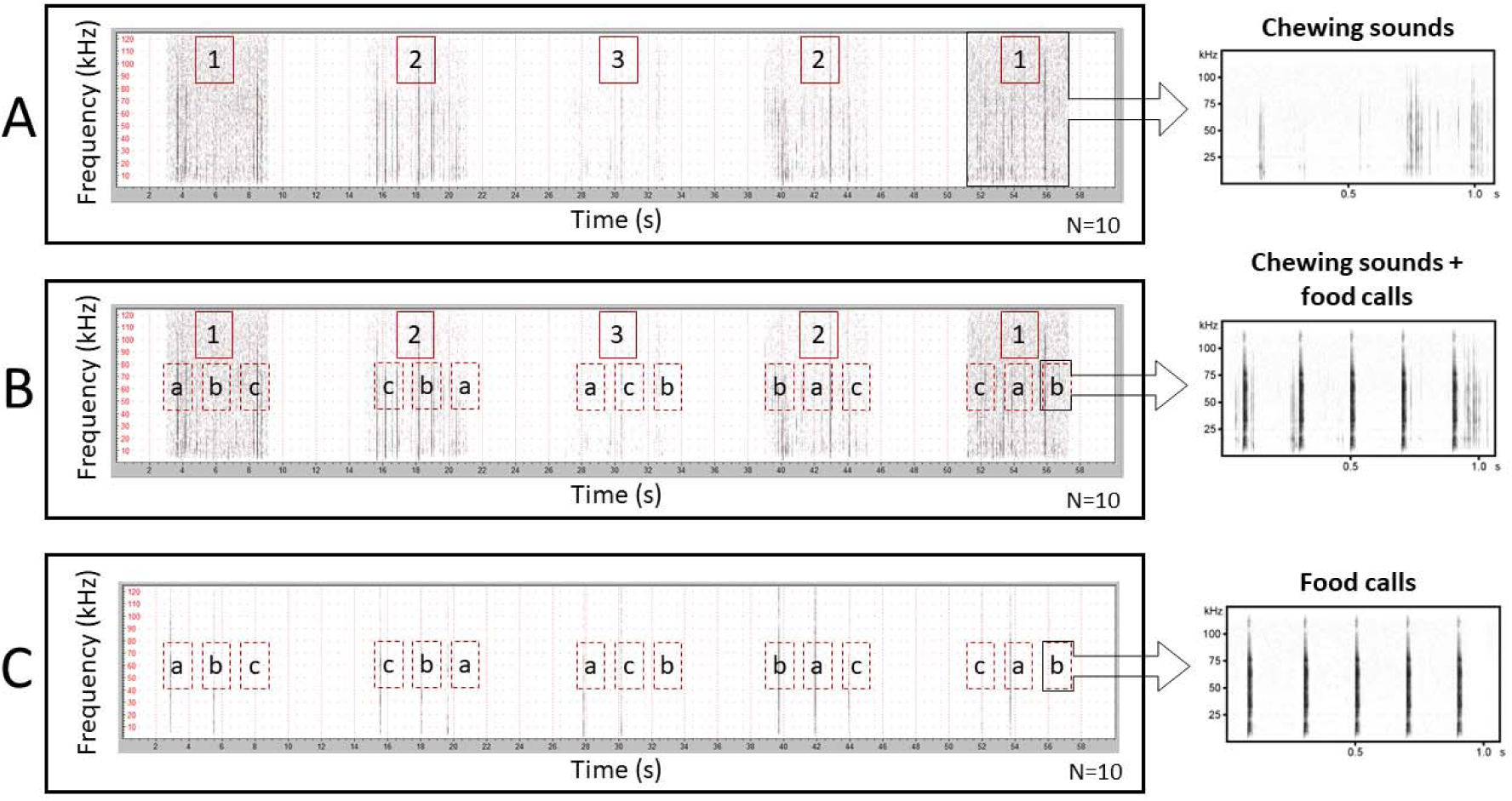
The setup of files used for playback experiments. A) First, we generated a 60 s wav file with 5 chewing sound bouts of 6 s each from 3 different individuals (shown in different numbers). In this example, we include two chewing sound bouts from individual 1, two bouts from individual 2, and one from individual 3. B) To the same file, A, we added 75 food calls, 15 per bout. Each call (e.g., c from the first bout) was emitted in bouts of 5, in rapid succession (time interval ≈ 0.20 s). The calls were obtained from 5 different individuals (e.g., a, b, c, d, e), and when possible, we used 3 different calls per individual. C) Finally, we removed the chewing sounds from the files used in B, keeping the 45 food calls with the same time intervals. A total of 10 files were created for each playback type (A, B, and C).

Using these 32 sound files (10 chewing sounds, 10 food calls, 10 chewing sounds + food calls, 1 pink noise, 1 silence), we generated 40 playlists in Avisoft Recorder USGH software. We randomized the order in which each type of sound was emitted and the sound file that was used for chewing sounds, food calls, and chewing sounds + food calls stimuli. In addition to these 5 files, each playlist also included a 15 s silence period before the start of experiments, then a short 0.5 s sound to indicate the start of playback. We also included a short 0.2 s sound to indicate the transition between stimuli and another to indicate the end of the playback trial.

### Experiment 1: Onset of eating

We investigated two possible effects of food calls on receivers: (1) a shift in the onset of eating novel prey (mealworms), and (2) a shift in exploratory or approach behavior. To test for the first effect, we captured naïve bats (adult individuals that never been fed mealworms) and broadcast food calls while holding them with a mealworm next to their mouths. For these experiments we individually placed bats at 10 cm from a speaker (Vifa, Avisoft Bioacoustics) connected to a USG Player (UltraSoundGate Player 116H, Avisoft Bioacoustics) and under a video camera (SONY FDR-AX53) using night-vision mode. All experiments were conducted under red light. We also placed a CM16 microphone directed towards the speaker and near the bat. The microphone was connected to an UltraSoundGate Recorder (model 116 Hme, Avisoft), and both the USG Player and Recorder were connected to a laptop computer running Avisoft Recorder software. The headphone outlet of the USG recorder was connected to the microphone inlet of the video camera, which allowed us to synchronize the video with the audio that was broadcast for the following analyses. We only include results for bats that fed during our playback trials (n = 15). We defined the onset of food consumption as the individual accepting and beginning to eat the offered mealworm. Videos were analyzed using the software BORIS (Friard & Gamba, 2016) to determine which stimulus prompted the onset of food consumption.

### Experiment 2: Approach and exploration

In the second experiment, we investigated whether the playback of food calls or chewing sounds would elicit exploratory or approach behaviors. Each focal bat was placed individually in a small square pyramid-shaped cage (1.5m long x 1.5 m wide, 1.5 m from base to apex) lined with mosquito net. Inside the cage we placed a plastic container (30 cm height), covered in cloth, in one corner of the flight cage. Next to the top of the container, and outside the cage, we placed a speaker (Vifa, Avisoft Bioacoustics). Next to the speaker we placed a CM16 microphone; this allowed us to synchronize the video with the audio that was being played back, as in the previous experiment. We traced two semicircles from the corner where the speaker was placed, the first one was placed at 50 cm from the corner, and the other at 75 cm.

Each bat was fed one mealworm shortly before the onset of the first trial. Trials started at 6:30 pm, when a single bat was released within the flight cage. We started playback upon the bat’s release; after the playback ended, we captured the bat using a hand net. All other bats were kept in an acoustically isolated location to avoid acoustic interference. Each bat was tested in three separate occasions, each time using a different playback list. At the end of each trial, we fed each bat a small mealworm and provided water *ad libitum*. At the end of the third trial, bats were fed mealworms *ad libitum*. Video recordings were analyzed in BORIS (Friard & Gamba, 2016) to determine the amount of time bats spent in the 75 cm area. We also counted the number of occasions in which individuals entered the 50 cm region. We did not estimate time spent within the 50 cm region given the fleeting nature of each visit.

### Data analyses

We compared the number of food calls produced by bats based on their sex, whether they were feeding on mealworm larvae for the first time (naïve = yes) or not (after bats had eaten mealworms on a previous night the bats were no longer naïve = no), and trial number (first, second or third). To do this, we first created a global model that could include additive effects of all predictor variables or up to 3-way interaction effects using the function glmer (family = poisson) in the R package lme4, with bat identity as the random variable. We then used the dredge function of the R package MuMIn to generate a table of models that include every possible combination of predictor variables. We compared models using the delta AIC, and kept the best models based on delta values < 5. For that subset of models, we applied the function sw in MuMIn, which provides an estimate of the relative importance of each predictor variable given the number of times that variable was included in the best models. To interpret the relationship between explanatory and response variables, we created a model in which we kept only the explanatory variables, and their interactions, that were present in at least 50% of the best models (i.e., those with delta AIC < 5), based on their relative importance. With this latter model, we determined which factors were significant (p value < 0.05). To test the effect of the retained fixed factors we used the anova function in the R package lmerTest (Kuznetsova et al., 2017). Only those factors that were significant were kept for post-hoc tests, using the function lsmeans with pairwise comparisons in the R package emmeans.

To extract spectral and temporal parameters of food calls, we used SASlab Pro (Avisoft Bioacoustics). We retained only the best quality signals, i.e. those with high signal-to-noise ratios. We manually labelled the calls and then measured peak, minimum and maximum frequencies, in addition to entropy, for the following positions: center of element, maximum amplitude of element, and mean spectrum of entire element. Spectrograms were analyzed using a an FFT length of 512 kHz (bandwidth resolution = 977 Hz), with an overlap of 93.75% (temporal resolution = 0.06 ms).

We analyzed our results of the first experiment by first comparing the number of times a given stimulus prompted the onset of eating during that trial. We first used a generalized linear model (function “glm” in the R package lme4; Bates et al., 2015) with a binomial distribution, with onset of eating as our response variable, where 1 was assigned to a stimulus in which feeding occurred for the first time, while 0’s represented either no feeding or feeding did not occur then for the first time. As explanatory variables we used stimulus type (silence, pink noise, chewing sounds, food calls, chewing sounds + food calls) and trial number (first, second, third, fourth or fifth) in an interaction model. For post-hoc tests we used the function “lsmeans” in the R package lsmeans (Lenth, 2016). For the latter we transformed data to percentage, and values of 0 (a stimulus in which the onset of eating never occurred) were increased to 1 (out of 100) to avoid errors in the estimate. We present results based on estimates of the function “pairs”.

In our second experiment we tested whether the amount of time spent within the 75 cm region, or number of times bats visited the 50 cm area, varied based on stimulus broadcast. For this analysis, we used time (model 1) and number of visits (model 2) as response variables and various explanatory variables, including stimulus, stimulus order (1 through 5), and trial number (either the first, second or third trial per individual). To address the possibility of a lingering effect from the previous stimulus on the tendency of bats to approach the speaker, we also included as a potential explanatory variable the stimulus broadcast immediately prior to the stimulus under analysis. For example, when analyzing the effect of pink noise on time spent in the 75 cm region, we determined whether food calls had been played back before this pink noise stimulus. We named these variables “pre-calls” and “pre-chew”. Because of the number of potential explanatory variables (n = 5), we created a global model that could include additive effects of all predictor variables or up to 4-way interaction effects using the function lmer (model 1) or glmer (model 2, family = poisson) in the R package lme4, with bat identity as the random variable. The selection of models, predictor variables, and post-hoc tests follow the methods explained above.

### Ethical Note

All sampling and experimental protocols followed guidelines approved by the American Society of Mammalogists for capture, handling and care of mammals (Sikes, 2016) and the ASAB/ABS’s “Guidelines for the Treatment of Animals in Behavioural Research and Teaching” (2012). This study was conducted in accordance with the standards of the Government of Panamá (Ministerio de Ambiente permit number: ARG-278-2022). Experimental protocols were also approved by the Smithsonian Tropical Research Institute’s Animal Care and Use Committee (IACUC protocol number: SI-23001).

In this study, we captured Spix’s disc-winged bats in the wild by searching furled leaves of various plant species in the order Zingiberales. To avoid disturbing bats while in their roosts, we approached the leaf very quietly and searched for bats with an extendable mirror; if the presence of bats was confirmed, we placed a transparent plastic bag at the opening of the leaf and carefully pinched the leaf at the bottom slowly moving towards the top, which caused the bats to crawl to the opening and into the plastic bag. Once the bats were in the plastic bag, they were transferred into cloth bags for transportation.

While performing flight cage experiments, we kept each social group together in the same bag to avoid social disturbance. Keeping individuals together like this does not result in conflict; it mimics natural social conditions, and appears to decrease stress. Moreover, we kept bags in a ventilated area with no direct exposure to sunlight. If bats were participating in individual trials, after the trial we returned them to the same bags.

At the end of the experiments, we provided mealworm larvae (*Tenebrio molitor*) and water to all individuals. We released the entire social group by placing all the individuals inside the same or a nearby leaf where they were found roosting earlier in the day. We have used this technique for returning bats to their habitat for over a decade, and individuals remain calm and in their natural roosting positions immediately after their return.

## RESULTS

We recorded a total of 15 bats from 8 different social groups as they were feeding on mealworms, to determine if they produced food calls. All individuals produced vocalizations except two, and bats typically emitted only a few calls per bout (Table 1). The results of model selection to determine how our explanatory variables influenced the number of calls produced show that the best models (2 models with delta values < 5) include the variables naïve (n = 2), sex (n = 2), and trial (n = 2) as additive effects, plus the interaction terms naïve*trial (n = 2), sex*trial (n = 2), and naïve*sex (n = 1; Supplementary Table S1). For post-hoc tests, the variable “sex” was dropped both as an additive term and from all its interactions, since it was non-significant. Our results show that naïve females in their first trial produced a significantly greater number of food calls compared to non-naïve females and males (Fig. 3). The model also predicts a larger number of calls from naïve males compared to those males that had consumed mealworms before, yet this comparison is based on results of only one naïve male bat.

**Fig. 3.**
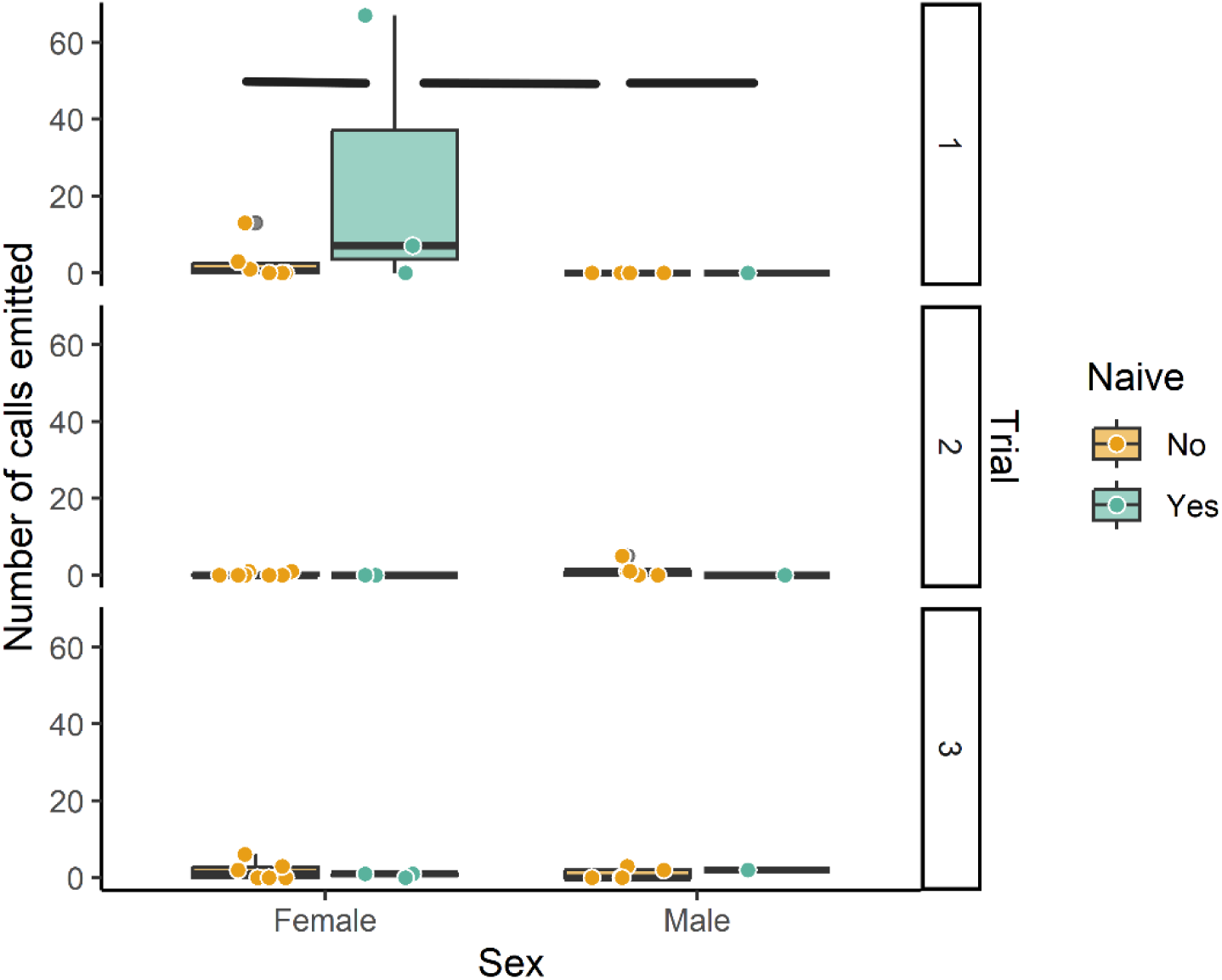
Number of food calls emitted by hand-held bats being offered mealworms. Results are divided by sex, trial number, and whether bats were feeding on mealworms for the first time (naïve = yes) or not (naïve = no). Horizontal lines represent significant (p ≤ 0.05) differences among naïve and non-naïve males and females based on the best model selection.

**Table 1.** Number of calls emitted by bats. Data are separated by trial number, sex, and whether bats were feeding on mealworms for the first time (naïve = yes) or not (naïve = no).

| Sex | Naïve | Bat ID | Number of calls |  |  |
| --- | --- | --- | --- | --- | --- |
|  |  |  | Trial 1 | Trial 2 | Trial 3 |
| Female | No | 982091068535013 | 0 | 1 | 0 |
|  |  | 982091068540751 | 13 | 1 | 6 |
|  |  | 982091068549575 | 0 | 0 | 2 |
|  |  | 982091068557450 | 1 | 0 | 0 |
|  |  | 982091068560202 | 0 | 0 | 3 |
|  |  | 982091068560731 | 3 | 0 | 0 |
|  | Yes | 982091068539737 | 67 | 0 | 1 |
|  |  | 982091068541705 | 7 | 0 | 1 |
|  |  | 982091068556544 | 0 | 0 | 0 |
| Male | No | 982091068539844 | 0 | 1 | 0 |
|  |  | 982091068543165 | 0 | 0 | 2 |
|  |  | 982091068550950 | 0 | 0 | 0 |
|  |  | 982091068551076 | 0 | 1 | 3 |
|  |  | 982091968552891 | 0 | 5 | 0 |
|  | Yes | 982091068548102 | 0 | 0 | 2 |

From the recording of these vocal individuals, we were able to obtain 19 calls of sufficiently good quality for analysis (i.e., those with high signal-to-noise ratios). Mean duration of calls was 0.017 s (standard deviation: ±0.005), with a mean peak frequency of 46 kHz (±15) at the maximum amplitude of the call. Minimum and maximum frequencies at the call’s maximum amplitude ranged between 15 and 82 kHz (±14). The average entropy of calls, at the call’s maximum amplitude, was 0.43 (±0.07). For further details and acoustic parameters see Supplementary Table S2.

We found that the stimulus that prompted the onset of eating was most often the food calls, either alone or in combination with chewing sounds (Fig. 4). Five bats started to consume the mealworms for the first time during the playback of calls, and 7 bats initiated consumption for the first time while hearing the chewing sounds plus calls playback. A few bats consumed mealworm larvae during the period of silence, some after hearing the playback of food calls (Supplementary Figure S1).

**Fig. 4.**
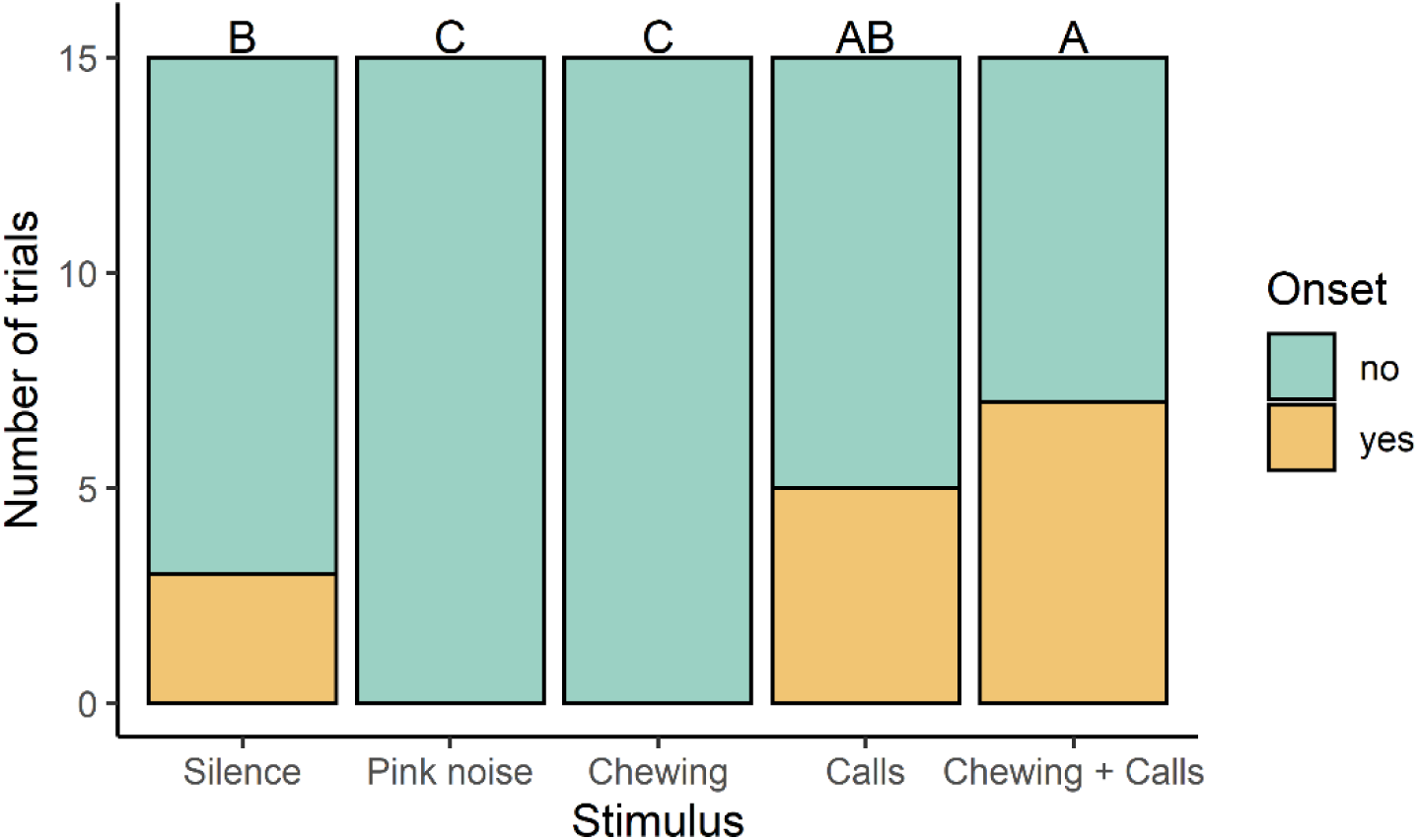
Number of trials in which a given stimulus prompted the onset of feeding. Different letters above bars denote significant differences among stimuli.

The results of model selection to determine how our explanatory variables influenced time spent near the 75 cm area around the speaker show that the best models (16 models with delta values < 5) primarily include the variables stimulus (n = 9), trial number (n = 16), pre-calls (n = 9) and stimulus order (n = 16) as additive effects (Supplementary Table S3). For post-hoc tests, the variable “pre-calls” was dropped as it was non-significant. The results of the model show that bats spent more time near the speaker during playback of the fourth stimulus, and bats spent increasingly more time near the speaker with each new trial (Supplementary Figure S2). We also observed that bats spent more time within the 75 cm area when food calls were being broadcast compared to the playback of pink noise (Fig. 5). However, time spent in the 75 cm area during the playback of food calls was not different than when silence, chewing sounds, and chewing sounds plus food calls were broadcast (Supplementary Figure S2).

**Fig. 5.**
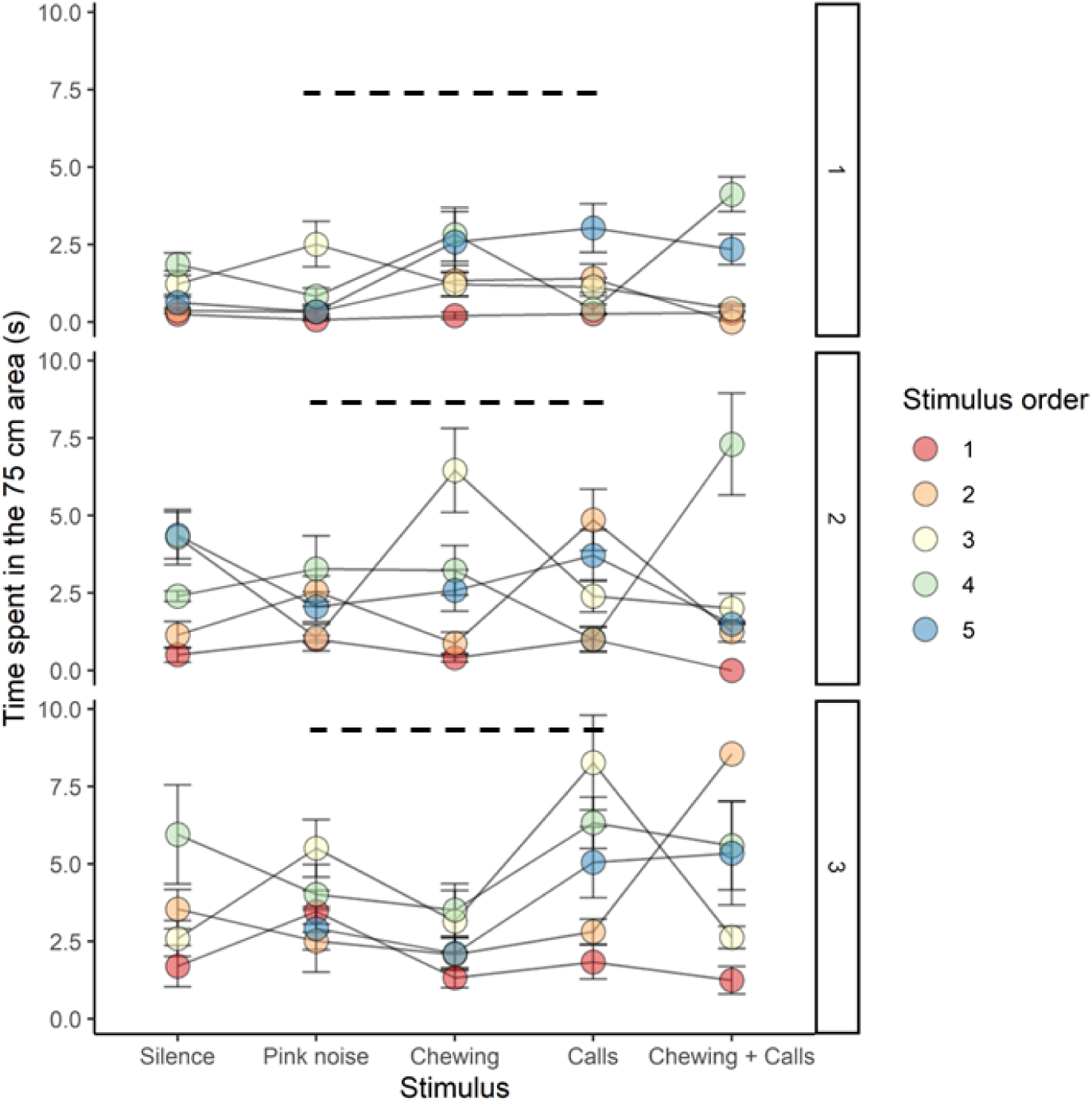
Time bats spent in the 75 cm region during each stimulus presentation. Data are arranged according to trial number (1, 2 or 3) and the stimulus order (1 through 5). Dashed lines represent borderline significant (p = 0.06-0.07) differences among stimuli. Circles represent mean values, and error lines standard errors.

The best models that predict the number of times bats visited the 50 cm area around the speaker (44 models with delta values < 5) primarily include the variables stimulus (n = 44), trial number (n = 44), pre-calls (n = 43), stimulus order (n = 33), and pre-chew (n = 29) as additive effects. However, other interaction terms were included in many (> 50%) of the best models, most notably pre-calls*stimulus (n = 32), pre-call*trial number (n = 32), and stimulus*trial number (n = 25; Supplementary Table S4). The variable “pre-chew” was dropped from further analyses as it was non-significant. All other additive and interaction effects were kept in the model for post-hoc tests. Results show that when calls had not been played-back before, bats approached the speaker more often during the playback of food calls compared to all other stimuli except pink noise, especially during the third trial (Fig. 6). Notably, while bats approached in response to food calls broadcast in isolation, the stimulus that included both chewing sounds and food calls did not prompt the approach of bats to the speaker.

**Fig. 6.**
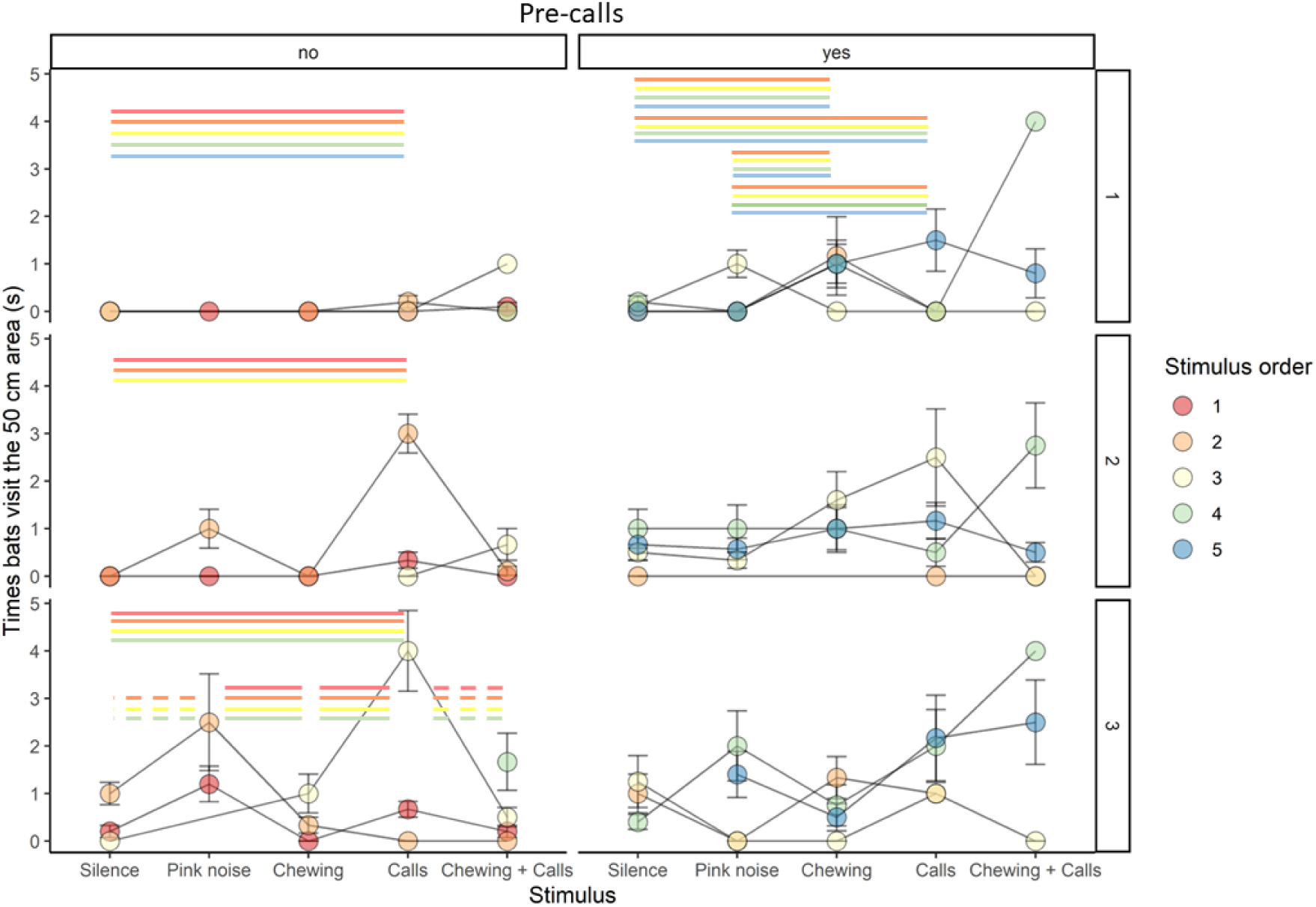
Number of times bats visited the 50 cm region during each stimulus presentation. Data are arranged according to trial number (rows: 1, 2 or 3), whether call stimuli had been presented before (columns: no, yes), and the stimulus order (1 through 5). Lines and dashed lines represent significant (p ≤ 0.05) and borderline significant (p = 0.06-0.07) differences, respectively, among stimuli based on the best model selection, with line colors denoting differences within the corresponding stimulus order. Circles represent mean values, and error lines represent standard errors.

## DISCUSSION

In this study we hypothesized that if the acoustic signals emitted by *T. tricolor* while feeding are indeed food calls, then their playback should elicit the consumption of provisioned food items in attending bats and the recruitment of conspecifics during flight. Our recordings of feeding bats show that many individuals emitted distinct vocalizations while consuming an abundant prey item. This type of call has not been previously described in disc-winged bats (Chaverri et al., 2018, 2021; Chaverri & Gillam, 2016; Montero & Gillam, 2015), and is markedly different from calls emitted by this species in situations of distress (Chaves-Ramírez et al., 2023). We also observed that vocalizations prompted two responses in receivers which are typically associated with food-calling (Clay et al., 2012), namely increasing feeding-related behaviors and social recruitment. Specifically, we observed that the onset of consumption of novel prey items was strongly associated with the emission of food calls but not with other types of sounds. Individuals also approached a speaker broadcasting food calls when flying in a tent, especially when calls had not been played-back before, while other types of sounds did not consistently prompt inspection. Taken together, these results strongly suggest that *T. tricolor* emits food-associated calls.

Food calling has been observed in several social species, including fowl (*G. gallus*), ravens (*Corvus corvax*), chimpanzees (*Pan troglodytes*), and white-faced capuchin monkeys (*Cebus capucinus*), among others (Clay et al., 2012). In these species, authors reported that calls are emitted exclusively in a feeding context, prompt social recruitment, may change with food characteristics, and can encode information about the characteristics of food, such as quality and quantity (Boinski & Campbell, 2010; Bugnyar et al., 2001; Gros-Louis, 2004a, 2004b, 2006; Hauser et al., 1993; Hauser & Wrangham, 1987; Heinrich & Marzluff, 1991; Marler et al., 1986; Slocombe & Zuberbühler, 2006). In bats, only one study to date has reported vocalizations that may be considered food-associated calls. Wilkinson & Boughman (1998) found that greater spear-nosed bats (*Phyllostomus hastatus*) emit “screech” calls while traveling to, and at, feeding sites, and primarily if individuals travel in groups. The emission of these calls prompts recruitment of conspecifics, both at feeding sites and also at the entrance of roosts (Wilkinson & Boughman, 1998). Therefore, the production of “screech” calls in this species, while strongly associated with foraging activities, is not considered to be exclusively associated with a feeding context (Clay et al., 2012). In contrast, the evidence collected in our study, together with results of previous studies (Chaverri et al., 2018, 2021; Chaverri & Gillam, 2016; Chaves-Ramírez et al., 2023; Montero & Gillam, 2015), shows that food-associated calls in *T. tricolor* are emitted exclusively in a feeding context, and that calls prompt social recruitment. Future studies are needed to confirm that *T. tricolor* emit food-associated calls while freely foraging, particularly in swarm-feeding situations, and to test whether these calls are modulated by, and may encode information about, specific food characteristics.

Food calling has only been observed in species that engage in social foraging. Social foraging is more likely to occur when food is abundant and clumped (Clay et al., 2012; Giraldeau & Caraco, 2000). In bats, social foraging is also associated with resource ephemerality (Kohles et al., 2022) and has been observed in species known to have strong social bonds (Egert-Berg et al., 2018; McCracken & Bradbury, 1981; Ripperger & Carter, 2021). *Thyroptera tricolor* lives in social groups with strong cohesion and high relatedness (Buchalski et al., 2014; Chaverri, 2010), and individuals often consume prey items found in large, yet ephemeral, insect swarms (Dechmann et al., 2006). Therefore, given its food-associated calling and combination of habits and traits, *T. tricolor* is an excellent contender for social foraging (Kohles et al., 2022). To provide further evidence of social foraging we not only need to track group members at night (e.g., Egert-Berg et al., 2018), but also explore in greater detail the diversity of prey items in *T. tricolor*’s diet using DNA metabarcoding (Sousa et al., 2019) to determine if food sources are typically found in abundant and clumped patches. Diet overlap within groups, and differences among groups, would provide further confirmation of social foraging (van der Post & Hogeweg, 2006).

While many bats in our study produced food-associated calls, we also found large inter-individual differences in the emission of these vocalizations. Specifically, we found that naïve females produced a greater number of food calls than males in their first trial. Individuals could be emitting a larger number of food calls in the first trial if food-associated calling reflects a signaler’s level of arousal in response to a feeding event (Caine et al., 1995). Females could be vocalizing more than males to help close kin locate a profitable food source. *Thyroptera tricolor* lives in matrilineal societies, exhibiting high levels of relatedness, particularly among females (Buchalski et al., 2014). Similarly, rhesus macaques (*Macaca mulatta*) live in matrilineal societies (Chepko-Sade & Sade, 1979), and females are known to produce more food-associated calls than males (Hauser & Marler, 1993). Despite the latter, more data are needed to understand the causes and consequences of the variation in the emission of food-associated calls within and among individuals. For example, individuals could change calling rates during the mating season. In *T. tricolor* we have seen sex-related differences in the emission rates of other vocalizations; for example, males typically produce a larger number of contact calls than females (Chaverri et al., 2020), especially during the mating season (Hernández-Pinsón et al., 2021). Calling rates could also be influenced by the audience (Dahlin et al., 2005) or by the type of food (Gros-Louis, 2004b; Hauser et al., 1993).

In conclusion, we found strong evidence that *T. tricolor* emits food-associated calls. While the primary advantage of food calling is conveying the location of food sources that can be shared among conspecifics, calling to recruit conspecifics can also protect the emitter from potential predators and intruders (Clay et al., 2012; Giraldeau & Caraco, 2000). Given that many species of bats are social (Kerth, 2008; McCracken & Wilkinson, 2000), that their food sources are often abundant and clumped (Kohles et al., 2022), and that social recruitment could potentially aid in the defense of these resources and provide protection against predators (Chaverri et al., 2018), it is puzzling that no other studies have provided evidence of food calling in this large mammalian order. Some of the main problems precluding further advances in this topic probably relate to the nocturnal habits of many species, their high mobility, and their small size. Direct observations of feeding behaviors and associated acoustic signals are often more feasible in diurnal species. Fowl and rhesus macaques, for example, can be followed on foot by researchers, enabling the direct pairing of observations between feeding and the emission of calls. Some species of bats can be fitted with GPS-microphone devices that provide great spatial resolution and the ability to record the vocalizations emitted throughout an individual’s foraging activities. Unfortunately, these devices are still too heavy for most species of bats, including *T. tricolor* (which weigh approximately 4 grams), precluding us from recording the calls emitted by individuals while foraging freely in nature. Further efforts to decrease the size of GPS-microphone tags, coupled with semi-captive observations and experiments, are needed to better understand food-associated calls, the context in which they are produced, and the information they convey in *T. tricolor* and other species of bats.

## Acknowledgements

We would like to thank Gregg Cohen for his insight, resourcefulness, and support during this project. We would also like to thank the many people that accompanied us during field work and experiments: Nair Cabezón, Eric de Framond-Bénard, María Sagot, Nicole Rose, Moisés Castillo, Grant Maslowski, Katie Galleta, Dyess Harp, and Jimena Víquez. GC would also like to thank the entire Smithsonian Bat Lab and administrative personnel at STRI, especially Adriana Bilgray, Aristoteles Villegas, Hilda Castañeda, Melissa Cano, Taisha Parris, and Orelis Arosemena for their valuable support.

## Author contributions

Conceptualization: G.C., R.A.P.; Data curation: G.C.; Formal analysis: G.C.; Funding acquisition: G.C., R.A.P.; Investigation: G.C., R.A.P.; Methodology: G.C., R.A.P.; Project administration: G.C.; Resources: G.C., R.A.P.; Visualization: G.C.; Writing - original draft: G.C.; Writing - review & editing: G.C., R.A.P.

## Data availability

A full data set and the code can be found at the GitHub platform: https://github.com/morceglo/Food-calling-Thyroptera.git.

